# Seeking voluntary passive movement in flies is play-like behavior

**DOI:** 10.1101/2023.08.03.551880

**Authors:** Tilman Triphan, Wolf Huetteroth

## Abstract

Play-like behaviour (PLB) is pervasive across the animal kingdom, especially in vertebrate species. Invertebrate PLB has been restricted to social or object interaction. Here we examined individual PLB in the vinegar fly *Drosophila melanogaster* by providing voluntary access to a spinning platform – a carousel. We demonstrate that flies exhibit idiosyncratic carousel interactions that qualify as play-like behaviour. While some flies show spontaneous avoidance, others actively seek stimulation, engaging in repeated, prolonged visits to the carousel. We propose that flies voluntarily expose themselves to external forces to intentionally receive exafferent stimulation. Self stimulation provides an efficient way to improve self-perception via internal model training and can shape multisensory integration.

**One-Sentence Summary:** Vinegar flies seek passive movement.

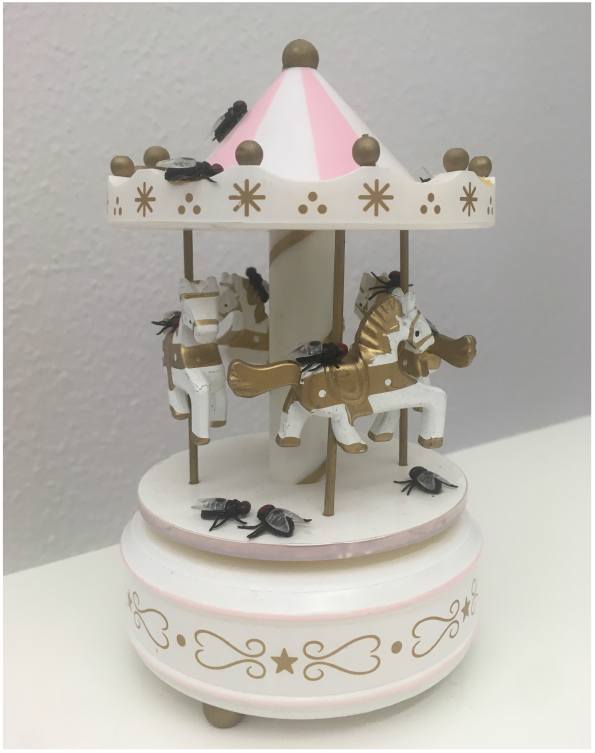

## Main Text

Anecdotal accounts about animals repeatedly exposing themselves to sources of passive movement by engaging with swings, slides, or carousels are frequently shared on social media. Generally, such behavior is assumed to be ‘play’, which, according to primatologist Phyllis Dolhinow, is *“(…) hard to define, but I know it when I see it (…)*” (*1*). More reliable criteria for play-like behavior (PLB) require the activity to be 1) of no immediate relevance for survival, but also 2) voluntary, intentional, rewarding, 3) non-ethotypical, 4) repeated, but not stereotyped, and 5) free from stress (*2, 3*). PLB following these rules has been found across the vertebrate subphylum (*3*); a recent study in rats even managed to identify involved brain regions (*4*). In invertebrates, the sparse reports so far addressed either social play in parasitoid wasps or spiders (*5, 6*) or object play in bumblebees (*7*). Contrary to the former PLB categories, a convincing hypothesis of the adaptive value of voluntary passive movement PLB is currently lacking. Flies are highly sensitive to the direction of gravitational pull (*8*), hence such intentional passive motion could be sufficient to externally induce proprioceptive stimulation (*9*). Here, we identify voluntary spinning on a carousel as idiosyncratic PLB in the vinegar fly *Drosophila melanogaster*, and propose how this behavior represents a highly efficient way to train internal body models.

## Results

We developed a behavioral apparatus to study voluntary passive PLB in flies, which consisted of an ‘enriched environment’, with access to food/water and a constantly spinning disk (a carousel, Fig. 1A). We exposed single flies to this experimental environment (ON) for several days and compared their behavior to that of flies exposed to the same arena with a stationary carousel (OFF). We restricted this study to male flies, given that they exhibit higher diurnal locomotor activity (*10, 11*) and a more conserved sleeping pattern (*12*), which mostly seemed to occur on the food patch (Fig. 1B, C). Experiments lasted up to two weeks (3-4 days on average) without any human interference, to provide a stress-free environment. For data comparability, and to allow for the novelty response to the unfamiliar environment to subside, we restricted our analysis to the first two complete consecutive days and nights. Both the spinning platform and the food spot were level with the rest of the arena so that negative geotaxis location preferences were negated (*8, 13*). Flies are capable of landmark and place learning (*14*) and can avoid unpleasant positions (*15*). Since we provided a high contrast landmark pattern around the arena, visits to the carousel’s position only depend on intrinsic motivational drive. Hence this behavioral paradigm is different from forced passive movement, which evokes stress-and even depression-like symptoms in flies (*16*).

**Fig. 1:**
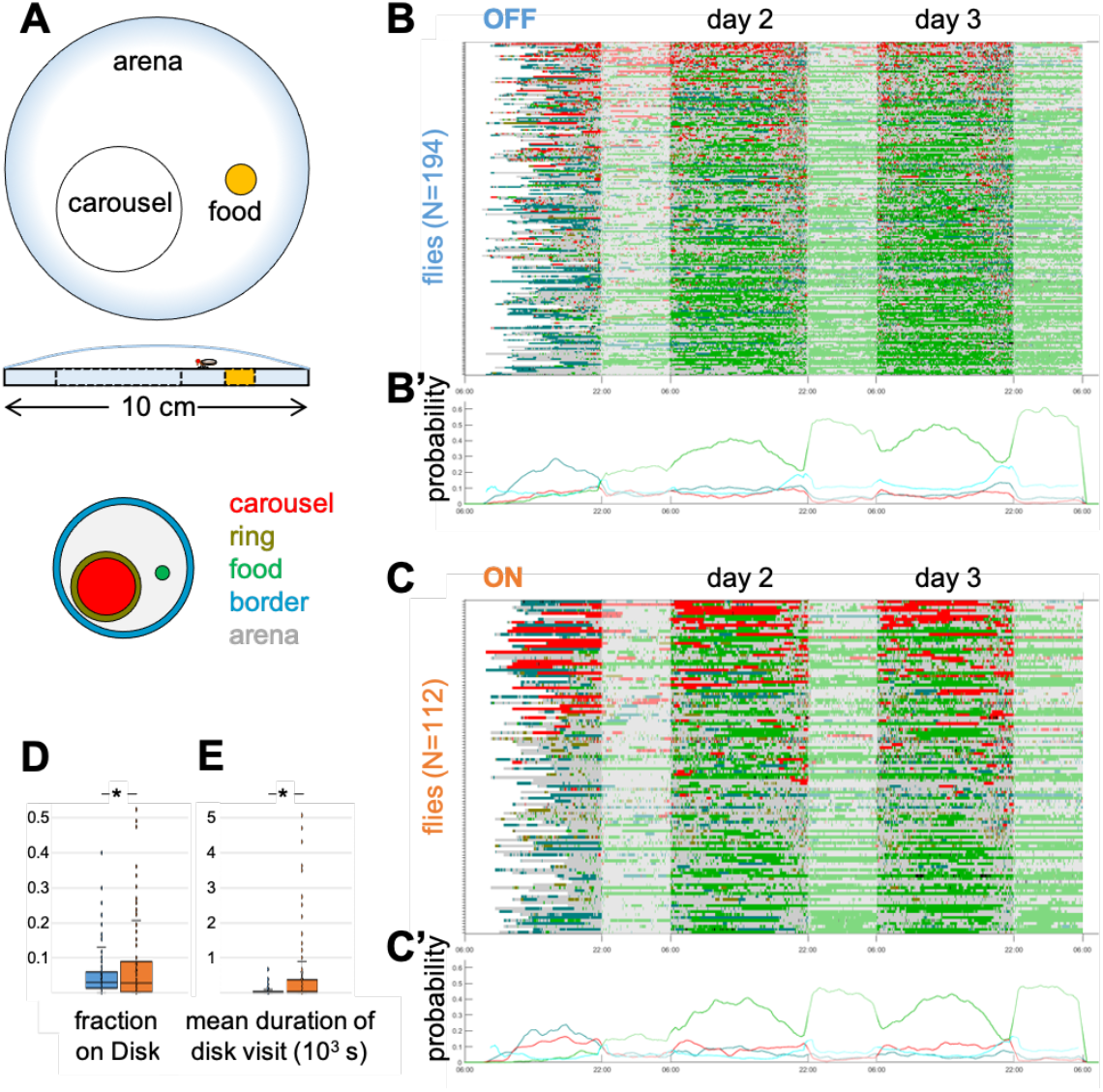
Carousel setup and idiosyncratic interaction. **A** The setup arena (10 cm diameter) contains a continuously spinning platform (4 cm diameter, 360°/s) and a food spot (1 cm diameter). The arena is covered by a Sigmacote-coated glass dome, a 16:8 LD rhythm is provided by IR and white light LEDs. Regions of interest (carousel, ring, food, border, arena) are defined for further analysis. **B, C** Experimental ethograms and **B’, C’** occupancy probability graphs (rolling window smoothing with 18.000 frames) over three nights (shaded) and two consecutive days of control flies (carousel OFF) and experimental flies (carousel ON). Elevated border exploration occurs at the beginning in both groups (teal). Circadian behavior settles from the second day onwards, e.g. increased food visits at night and during mid-day (green) and rhythmic activity (cyan). The ‘ring’ ROI was omitted from occupancy probability graphs for clarity reasons. Carousel occupancies in experimental flies are more biased towards either less fragmented and prolonged visits or avoidance, mostly during day periods (red). **D** ON flies spend significantly more time on the carousel than OFF controls. Two-tailed t-test, p < 0,01. **E** Individual mean carousel visits are significantly longer in ON flies as compared to OFF controls, with a massive increase in variance (s^2^OFF = 1,8×10^5^, s^2^ON = 2,7×10^7^). Two-tailed t-test, p < 0,001. Data in D & E originates from clipped 48 h, beginning with day 2.

After an initial period of increased arena boundary exploration, both OFF control flies and ON experimental flies exhibited pronounced idiosyncratic place preference and locomotion, which followed overall diurnal activity (Fig. 1B, B’, C, C’). As reported for idiosyncratic behavior (*17*– *22*), these individual patterns were conserved across days. Initial boundary exploration and diurnal activity patterns disappeared in both OFF and ON groups after randomly shuffling the order of individual region occupancies, which favors the interpretation that these observed patterns in the experimental data are not an artifact (Fig. S1). Temperature and humidity had a significant impact on general fly activity (Fig. S2) as reported previously (*23*), but not on place choice (Fig. S3).

A crucial criterium of PLB is its voluntariness; here, ON flies were always able to access and leave the moving disk on their own volition. Interestingly, the overall length of time flies spent on the carousel was significantly higher when the carousel was spinning compared to when it was not (Fig 1B, C, D). The duration of the average carousel visit also increased significantly, with a large increase in variance (Fig. 1E). Some ON flies chose to avoid the platform or spent long periods of time immediately in front of it. Since flies are known to respond to an unexpected, potentially threatening cue with evasive responses (*24*–*26*), this could explain carousel avoidance, but not voluntary, repeated exposure.

To further understand the idiosyncratic interactions the flies exhibited with the moving disk, we selected the ON flies with the highest probability of spending time on the carousel (>5% of time, ‘seeker’), and those with the lowest probability of spending time on the carousel (up to 0.5% of time, ‘avoider’). The environmental occupancy of these groups of ON flies differed substantially from each other and from control OFF flies (Fig. 2A-C). All groups spent most time on the food, but while OFF control flies showed considerable border exploration and no obvious preference or avoidance for the carousel ROI, ‘seeker’ and ‘avoider’ were more drawn to the carousel or the food, respectively. We next explored how often the flies in these groups transition from the carousel, in relation to the time spent on the carousel ROI (Fig. 2A’-C’). As expected, seeker flies exhibited more and longer carousel visits compared to controls, whereas avoider flies showed few carousel visits of shorter durations. Interestingly, we observed differences in behavioral choices following carousel visits (Fig. 2A’’-C’’). We assessed which regions of interest (ROI) were frequented within 100 s after defined lengths of carousel visits. For the OFF control condition, ROI visit probabilities were roughly aligned with relative ROI size, independent of preceding disk visit durations. Carousel-seeking ON flies however kept returning to the moving disk at increasing rates (Fig. 2B’’). In contrast, flies that mostly avoided the carousel did so consistently after a few visits and instead developed a subsequent preference for the border region (Fig. 2C’’). Both OFF control and seeker flies showed an increasing preference for food visits the longer they stayed on the carousel ROI, in line with work on hunger-motivated food exploration (*27, 28*). Hunger drive is in antagonistic competition with other motivational drives like the positively evaluated possibility to mate (*25, 29*), whereas thigmotactic border explorations are associated with fearful behavior in open field tests in flies, mice and humans (*30*–*32*). We therefore conclude that the interaction with a carousel enhances behavioral contrast between groups, suggesting idiosyncratic positive or negative association with this stimulus.

**Fig. 2:**
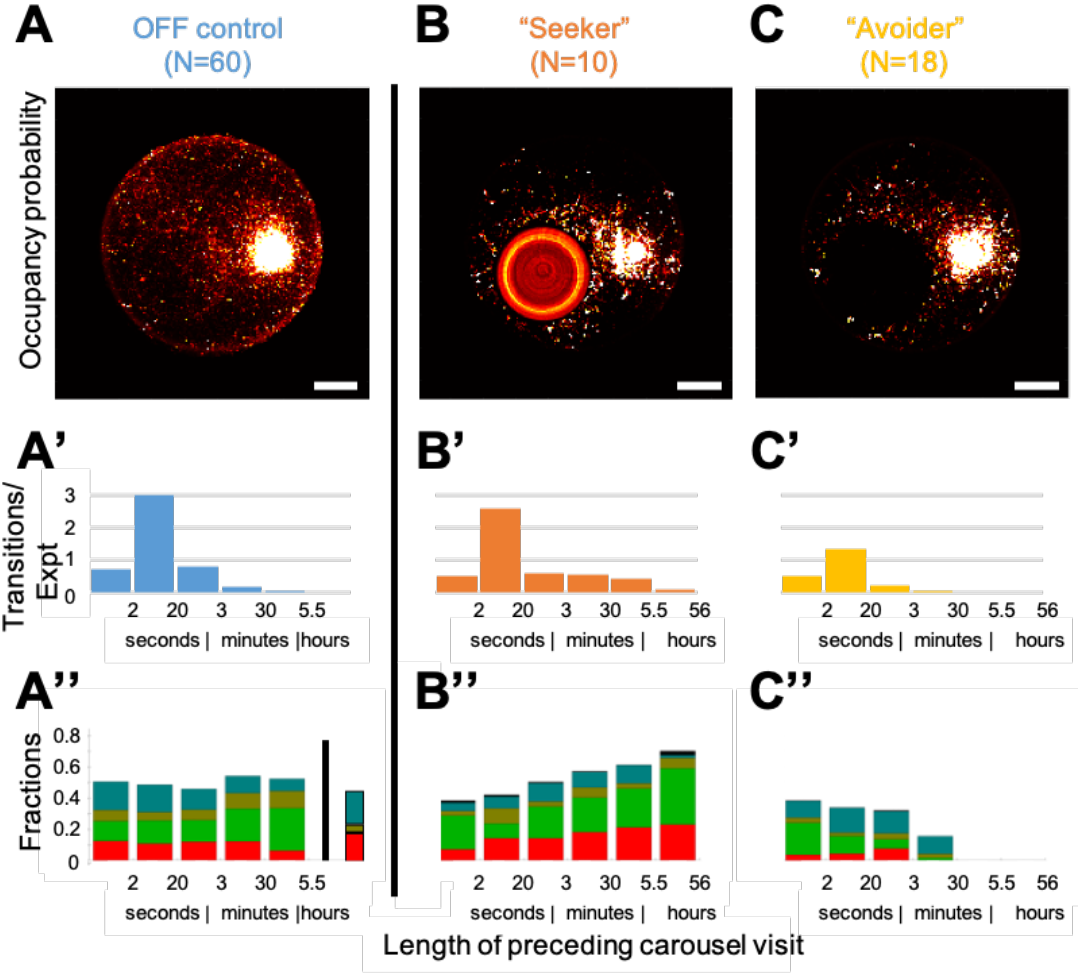
Carousel visits have divergent effects on subsequent behavior in idiosyncratically sorted fly groups. **A** Occupancy preferences in OFF control flies reveal preferred visits to the food and border region, while the stationary carousel remains inconspicuous in comparison to the remaining arena. Scale bar in A-C: 2 cm. **A’** OFF control flies rarely spent more than 3 minutes on the carousel ROI. **A’’** Stationary carousel stays of varying lengths have no major influence on subsequent arena region visits, except for an increasing preference for food visits. The leftmost stacked bar represents ROI fractions according to area size for reference (disk, red; food, green; ring, olive; border, teal). **B-B’’** Carousel-seeking flies exhibit an increasing probability of recurring carousel visits, dependent on length of stay, mostly at the expense of visits to the remaining arena. **C-C’’** Carousel-avoiding flies show the highest food occupancy. They rarely return to the carousel after accidental < 30 min visits, which are followed by increased border region visits. All N are selected for <0,005% unidentified positions/frame (NaNs, black fraction in B’’).

Finally, we addressed whether flies develop a positive association with voluntary passive movement. To do so we employed a spatial alteration paradigm with two carousels, spinning alternatingly for 5 minutes (Fig. 3A). We hypothesized that if flies positively value the exposure to voluntary passive movement they would be more likely to remain on the moving carousel until it stops. Once again, we ranked flies according to their disk occupancy and found that the 33 flies that spent the most time on the moving disks remained significantly more on the disk until it became stationary as compared to transitioning from a stationary to a moving disk (Fig. S4). In line with our previous results, this suggests that the carousel movement was not perceived as aversive.

**Fig. 3:**
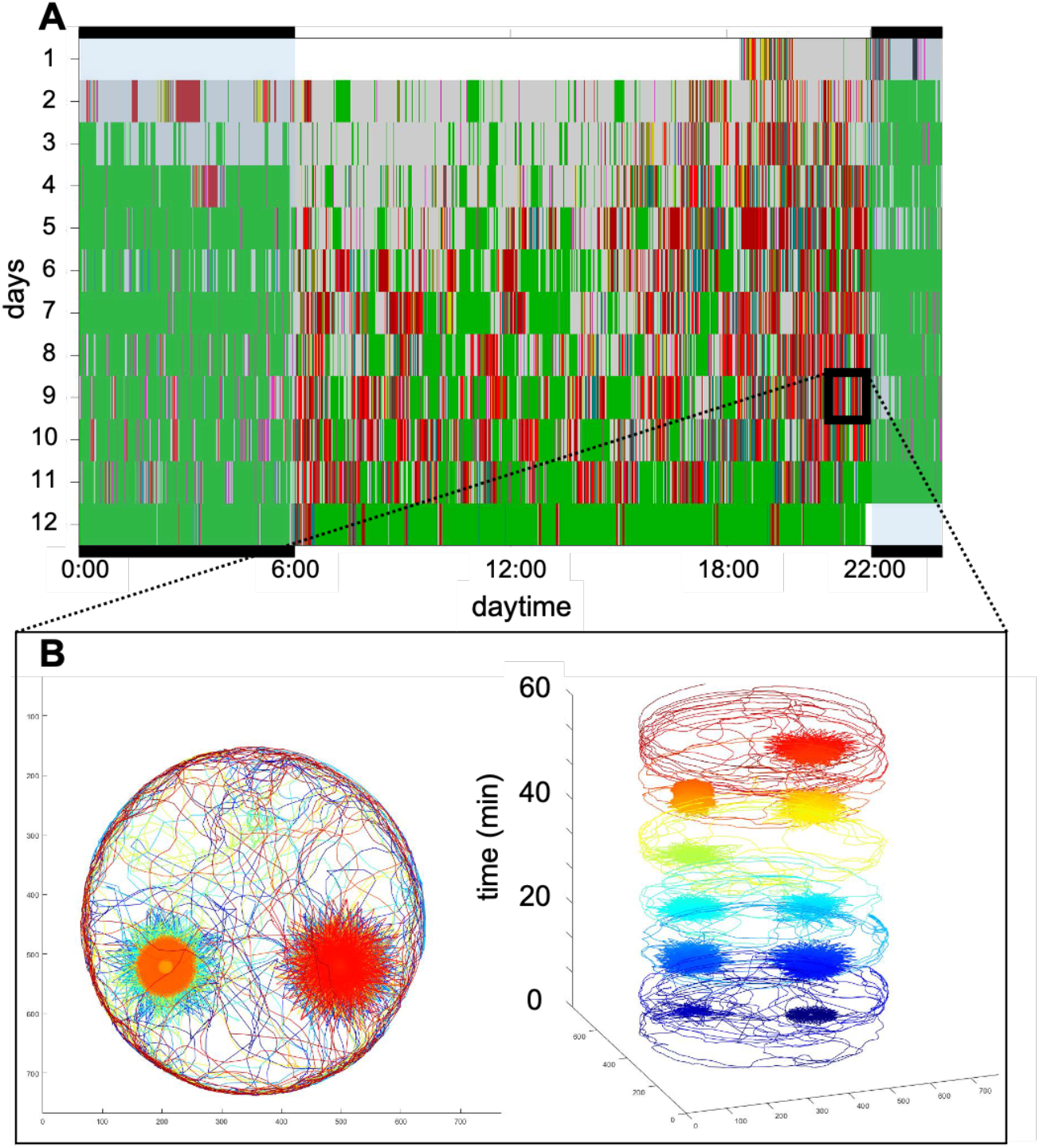
Flies seek spinning carousels. **A** Representative experimental ethogram for a single flies’ behavior over 12 days in a double carousel setup. Carousel visits preferentially occur during the evening activity peak and show no obvious pattern across days. **B** Walking traces during 1 h of the evening activity peak of day 9 reveal repeated, alternating visits to the respective moving carousel.

Although we cannot formally rule out that the flies felt trapped once they were on a moving disk, induced by the sensory experience of the visual flow or the centripetal force, the frequent revisits in the single carousel experiments (Fig. 2B’’) render this possibility unlikely. We nevertheless further investigated carousel seeking by testing whether flies alternated between the two carousels in the double arena. Such behavior would imply not only the active decision to go on a moving object, but to actively and repeatedly follow exposure to that stimulation. Indeed, such alternated sequences ‘tracking’ the ON carousel occurred at a significantly different rate to transitions between stationary carousels in an idiosyncratic fashion (Fig. S5). We even observed repetitive switching (Fig. 3B), strongly suggesting that the moving platform confers a situation-dependent positive value for individual flies -a key criterium of PLB.

## Discussion

In this study we established a behavioral paradigm with a carousel as a means of studying voluntary passive movement, which we assume might simulate a more natural setting of voluntary passive movement like sitting on swinging branches or leaves. Our data show that the flies’ active exposure to the carousel fulfils all criteria of play-like behavior (PLB): it is of no immediate relevance for survival, the flies engage with the carousel voluntarily, repeatedly, in a stress-free environment and in a non-ethotypical manner, and the seeking behavior strongly implies a rewarding component of intentional visits. The novel identification of PLB in this insect now allows for further analysis of the neuronal underpinnings of this curious behavior, utilizing the full genetic arsenal available in *Drosophila*.

But what could this PLB be good for? Some flies could seek the environmental enrichment and/or the sensory stimulation like individuals of many other species, which would allow for activity-dependent circuit refinement and would counter the negative effects of sensory deprivation (*33*). We hypothesize that voluntary passive movement PLB is particularly suited to refine neuronal circuits for improved proprioceptive acuity and self-representation. Exafference without creating an efference copy is achieved willingly; hence the corollary discharge of the internal model representation of the outcome of the planned action predicts the exafferent stimulus (*34*). This internal prediction can direct the incoming activity into efficient refinement of internal models of self-perception and, as derivative, lead to improved vision compensation and generally improved sensorimotor control and intersensory coordination (*35*–*43*). ‘Gamifying’, and hence rewarding this training of internal models as in PLB might be one way to motivate organisms to engage in such tasks that improve self-perception. Ultimately, a realistic representation of one’s own body (an embodied self) is a prerequisite for - at least - any bilaterian to efficiently and intentionally interact with the world (*44*).

We conclude that fly behavior exhibits insightful parallels to vertebrate action selection, which, in combination with previously demonstrated parallels in reward association (*45*) further supports the notion to pursue fundamental principles in neural circuit function using this accessible organism (*56*). The sustained boost in connectomic data availability for both the whole brain (*46, 47*) and premotor circuits of the fly (*48*–*50*) will allow us to pinpoint these computational principles to identified cellular components (*51*), and to assess how internal models are neuronally implemented (*52*).

## Acknowledgments

We thank Beate Dika, Christine Dittrich, Alexander Hergett, Clemens Sauter, Varvara Kenti Kranidioti and Yanan Zhang for help with generating the data. Clemens Sauter, Merlin Szymanski and Alex Hergett helped writing initial versions of software. Christine el Jundi, Clara Ferreira, Giovanni Galizia, Christoph Kleineidam, Carlotta Martelli, Randolf Menzel, Andreas Thum and Scott Waddell gave important feedback. Thomas Trenker, Fabian Rapparlie (both Konstanz) and Ingo Kannetzky (Leipzig) provided expert technical assistance.

## Funding

Zukunftskolleg Marie-Curie Incoming 2-year postdoctoral fellowship (TT); Zukunftskolleg Marie-Curie Incoming 2-year postdoctoral fellowship (WH); German Research Foundation grant HU 2474/1-1 (WH); German Research Foundation grant HU 2474/1-3 (WH)

## Author contributions

W.H. conceived the study, designed and supervised the experiments, acquired the funding and wrote the manuscript. W.H. and T.T. performed all data analysis, visualization and edited the manuscript. T.T. wrote and supervised all MATLAB analysis and Python recording software.

## Competing interests

Authors declare that they have no competing interests.

## Data and materials availability

All data are available in the main text or the supplementary materials.

## Supplementary Materials

## Materials and Methods

### Animals

CantonS animals ranged between 0 and 7 days in age. Flies were reared at 20-25°C and kept on standard fly food right before transfer into carousel arenas. Temperature and humidity were noted at the beginning of all experiments. Animal tracking lasted up to 346 hours or 2 weeks; most experiments were stopped after 3-4 days.

### Carousel layout

The setup consists of a 10 cm diameter arena, an off-centered 4 cm diameter spinning platform and an off-centered circular food spot (1 cm diameter, Fig. 1). Both spinning platform and food spot were levelled with the rest of the arena to avoid negative geotaxis location preferences (*13*). The arena was covered by a Sigmacote-coated (Sigma-Aldrich SL2) glass dome to deter roof walking and lit from above by 12 white or 12 infrared (800 nm) LEDs in a 16:8 LD rhythm. To provide visual landmarks, the outer walls were covered by randomly patterned black/white squares.

### Raspberry Pi recordings

A custom-written Python script automatically recorded batches of 30 min long videos with a resolution of 768×768 pixels at 5 fps as .h264 files, starting at the next half hour until manually interrupted. Lighting automatically changes to infrared LED light at 22:00 and back to white LED light at 6:00. Metadata (humidity, temperature, genotype, age, sex, food, date, time, additional information) is written into a .csv or .json datafile at the start of the experiment.

### Data conversion and preparation

Raw .h264 files are copied from Raspberry Pis onto an analysis PC with a custom-written program that automatically converts .h264 files into .mp4 files.

### Tracking

The fly position is assessed using DeepLabCut (*53*), which was manually customized to allow for batch processing. The network was trained on 10 frames from 40 flies each for 1,030,000 iterations, after marking the anterior and posterior end of the fly (or its center of gravity in later DeepLabCut versions) in each of these frames.

### MATLAB analysis

Tracking data in the form of .h5 files and corresponding metadata are batch-loaded into MATLAB using a custom-written software. All subsequent analyses are performed using an expanded and improved version of code already described before (*28*). Only experiments with less than 5% undetected fly positions/frame (NaNs) were used in the analysis (306 of 349). To consolidate data consistency, we restricted further analysis to day/night 2 and 3, after initial boundary exploration settled (303 expts with less than 5% NaNs). Some analyses required a more rigid threshold for NaNs (88 expts with less than 0.005% NaNs). All MATLAB code is available online on Github (https://github.com/triphant/DrosoCarousel).

## Notes

### Competing Interest Statement

The authors have declared no competing interest.

### Summary of Updates

Minor edits and streamlined discussion

