## Supplemental Figures 1-5 for "Seeking voluntary passive movement in flies is play-like behavior"

**Figure S1**

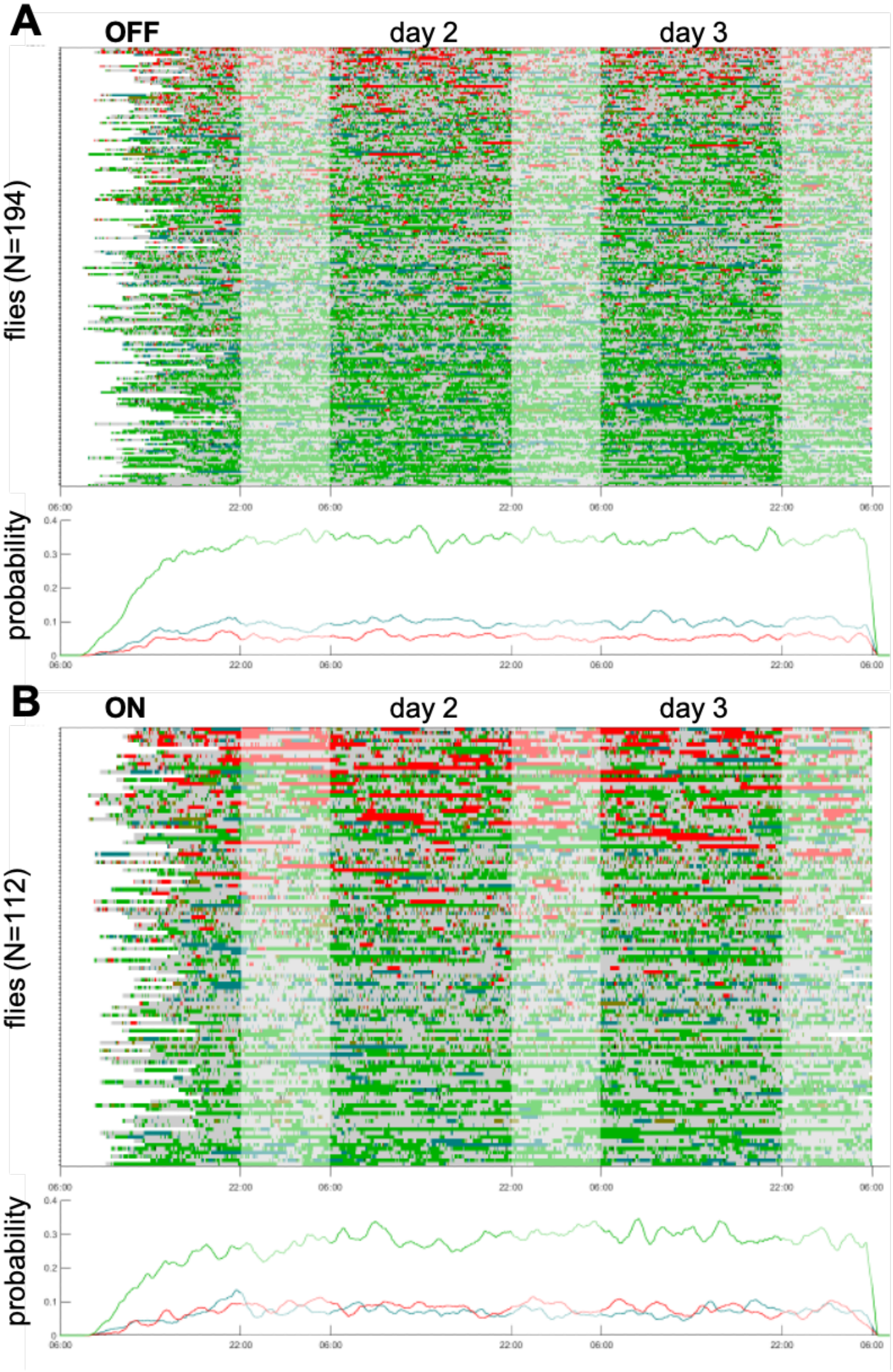


**Fig. S1: Behavioral pattern disappear in shuffled ethogram data**

Experimental ethograms of **A** OFF and **B** ON flies with shuffled events. Initial boundary exploration and diurnal occupancy probability pattern are no longer visible.

**Figure S2**

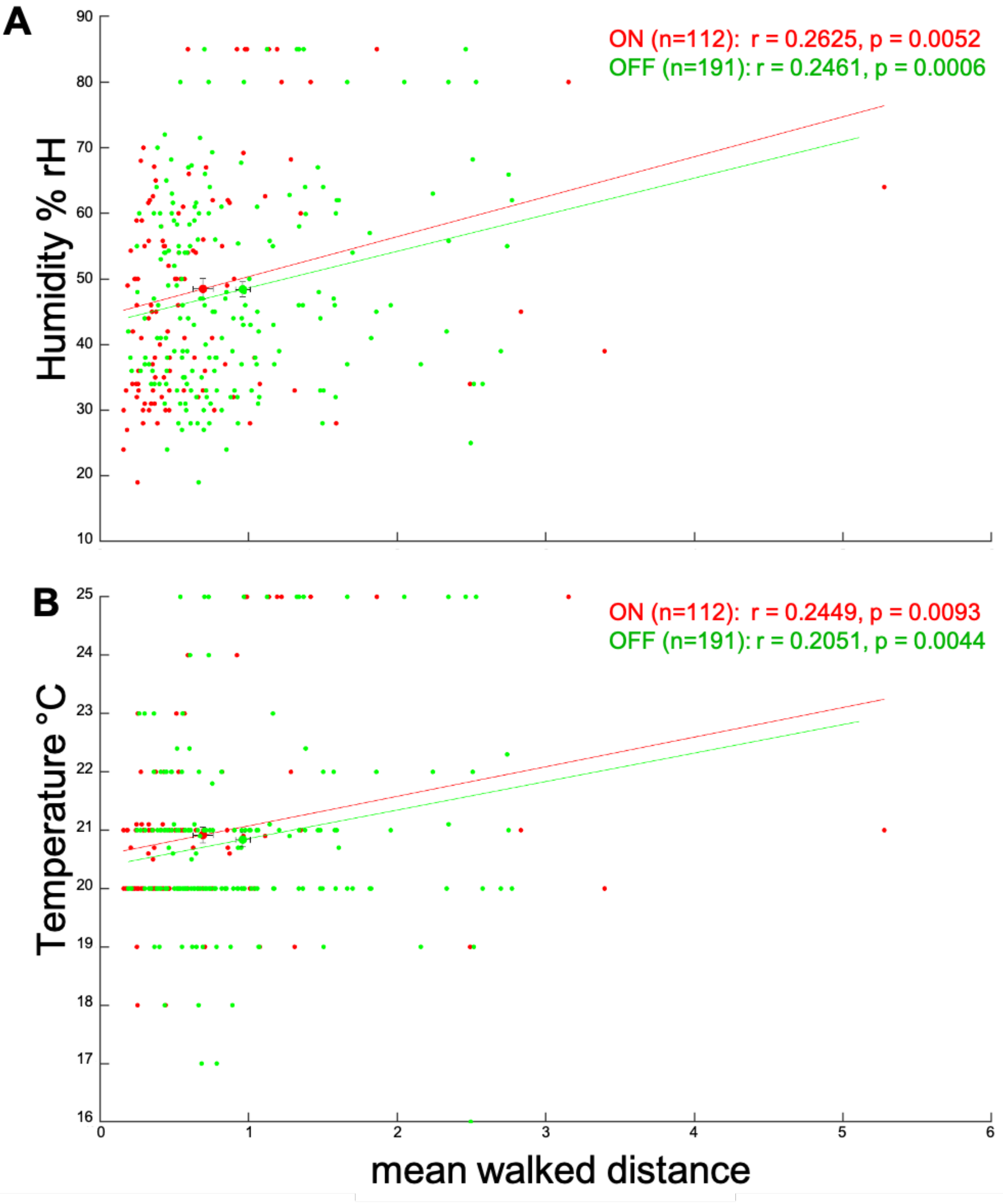


**Fig. S2: Humidity and temperature weakly influence activity levels**

**A** Humidity levels between 20-85 % rH and **B** temperatures between 17-25°C exhibit a positive correlation with activity levels (Pearson's correlation) in OFF and ON flies.

**Figure S3**

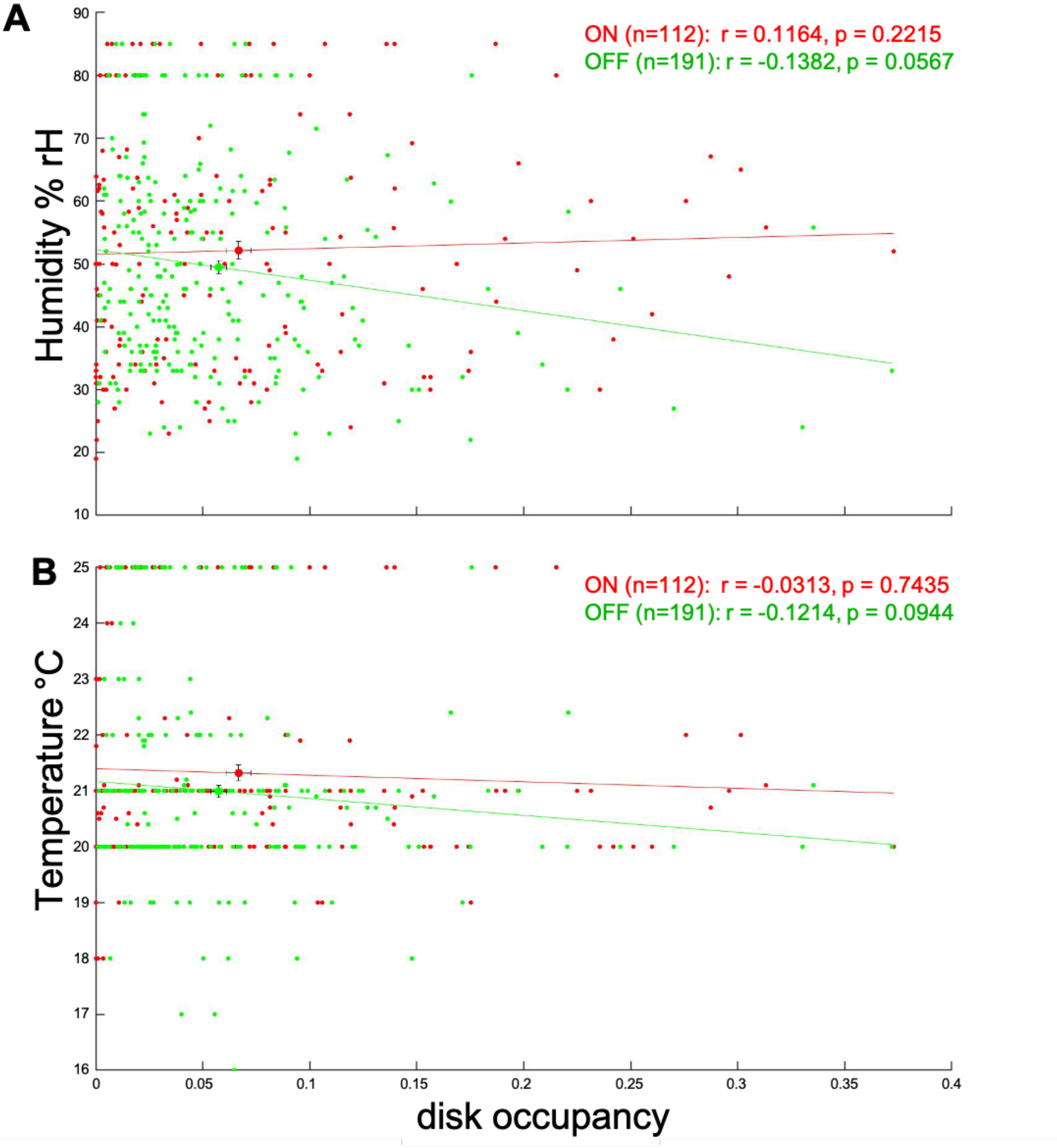


**Fig. S3: Humidity and temperature do not impact disk occupancy**

**A** Humidity levels between 20-85 % rH and **B** temperatures between 17-25°C have no impact on disk occupancy in OFF and ON flies.

**Figure S4**

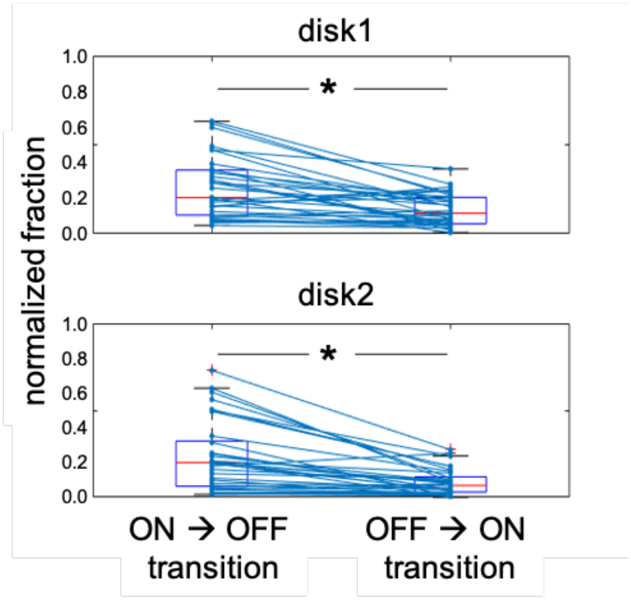


**Fig. S4: Flies seeking the moving disk preferentially stay until it stops**

The 33 "Seeker" flies in the spatial alteration experiments which spend the most time on moving disks are significantly more likely to stay until the end of movement (ON → OFF transition) rather than being trapped on a disk that suddenly starts to move (OFF → ON transition). Paired t-test, $p<0.0001$ for both disk1 and disk2.

**Figure S5**

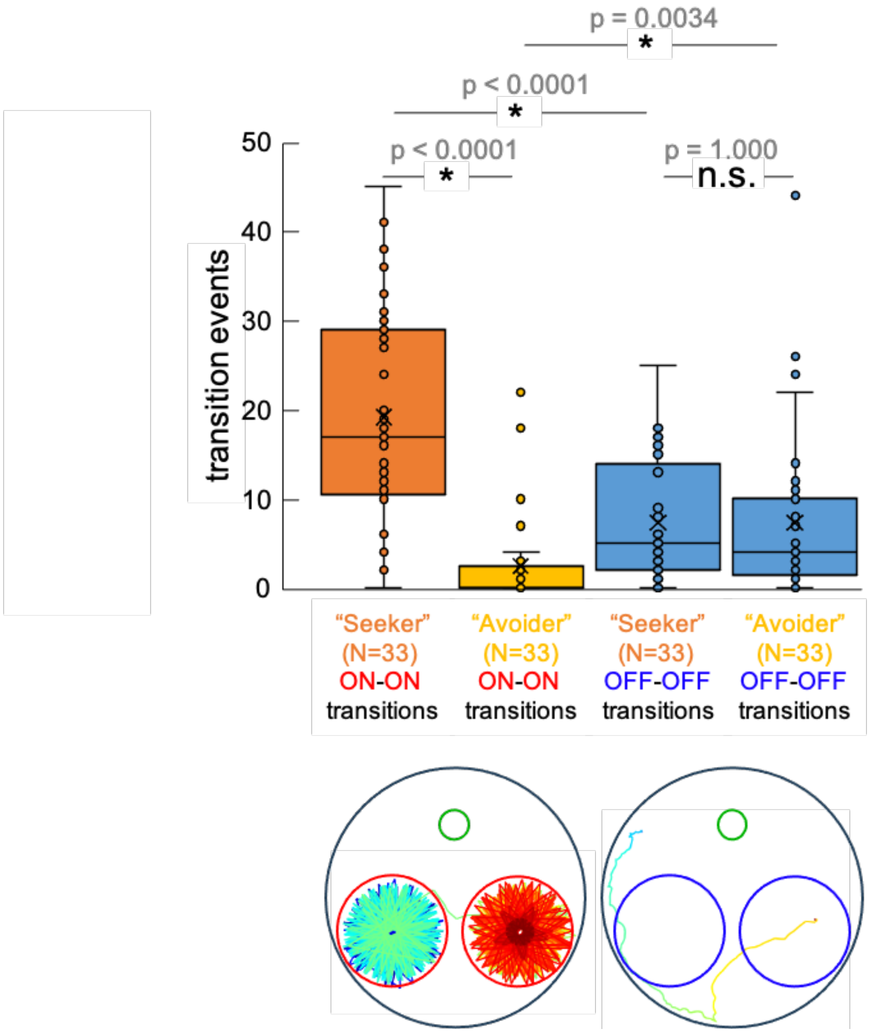


**Fig. S5: Idiosyncratic seeking of a spatially alternating moving disk**

The flies in the spatial alteration experiments (N=66) were ranked according to time spent on moving disks and then split in half to create "Seeker" and "Avoider" groups (N=33 each). Transitions from one moving disk onto the other within 100 frames were counted for both groups (ON-ON transitions). As reference, stationary disk transitions were also counted (OFF-OFF transitions). While the stationary transitions did not significantly differ between "Seeker" and "Avoider" groups, moving disk transitions occurred either significantly more ("Seeker") or less ("Avoider") frequently than corresponding stationary transitions. Paired or unpaired Student's t-test, respectively.
